# High Frequency of Dynamic Rearrangements In Crispr loci

**DOI:** 10.1101/2022.05.19.492656

**Authors:** Yaqing Ou, James O. McInerney

## Abstract

CRISPR-Cas immunization of prokaryotes proceeds by the acquisition of short fragments of invading DNA and integrating them into specific positions within the host genome in a process called adaptation. Adaptation is thought to be polarised, which suggests that CRISPR array spacer order reflects the recentness of the infection. The detailed processes through which CRISPR loci arise, and how they evolve are not completely clear. In this study, we collected 12,461 prokaryotic genomes, and using a combination of four different approaches and a series of conservative filters, we identified CRISPR arrays in 82.7% of Archaea and 40.6% of Bacteria. To understand spacer evolution in these CRISPR loci we firstly tracked point mutations in CRISPR repeats, and secondly, we carried out a comparative analysis of arrays that share multiple similar spacers. Both results indicate that CRISPR arrays are frequently dynamically rearranged. These findings are at odds with a model that suggests that spacer order is likely to reflect the recentness of infection. We conclude that the order of spacers in a CRISPR array, as well as the spacer content of the array, is likely to arise from a combination of events, such as insertion in the middle of the array, recombination within or between arrays, or Horizontal transfer of all or part of an array. We suggest these rearrangements are favoured by natural selection in complex and dynamic environments.

## Introduction

Approximately 40% of Bacteria and 80% of Archaea use CRISPR-Cas as a defence against invading prophage or other mobile genetic elements (MGEs) (1). CRISPR-Cas works by integrating modified genetic segments called protospacers into arrays on its host genome. Protospacers are acquired from viruses or MGEs (2), and once integrated, are called spacers (3). The entire process is known as adaptation, integration or acquisition. The result is a library of short fragments of foreign DNA, that are separated by highly conserved direct repeats. Upon subsequent infection, a spacer and part of its associated repeat can be transcribed and matured as CRISPR RNA (crRNA), which integrates and guides adjacent assembled Cas nucleases to destroy complementary invading DNAs (4). However, protospacers are selected in different ways during adaptation and the process depends on the type of CRISPR-Cas system. In type I, II and V CRISPR-Cas systems, the protospacer is only sampled near protospacer adjacent motifs (PAM) in the viral genome (5). PAMs can be thought of as a marker in the invader that the Cas protein recognised during adaptation. PAMs also help Cas proteins distinguish between self and invaders and avoid autoimmunity during interference (1, 6).

During CRISPR subtype I-E spacer adaptation, the Cas1-Cas2 complex needs to recognise the boundary between the leading sequence in a CRISPR array and the first repeat (7). This activity is dependent on an Integration Host Factor (IHF) that combines with the leading AT-rich segments and forms a U-shaped structure for recognition (8–10). In subtype II-A, there is a highly conserved leading-anchor sequence (LAS) located upstream of the first repeat, that ensures all newly integrated spacers are inserted at the leading side (Figure 1B) (11). In addition, the unique integration of foreign DNA, apart from naïve acquisition, can be triggered by an existing spacer in a CRISPR locus, and this process is called primed spacer acquisition (2, 12). When a prokaryote encounters a phage that contains a perfect or partial match to a spacer region, one or more additional protospacers can be acquired through primed acquisition upon interference (13). This process has been confirmed in both type I and type II CRISPR-Cas systems *in vitro* and *in silico* (14–17). However, both known integration paths are thought to add a spacer specifically to the leading side of the array (12). In other words, the order of spacers in CRISPR arrays is suggested to reflect the chronological order in which integration has occurred (18). McGinn & Marraffini (11) contended that the order of spacers in any CRISPR locus is a kind of “molecular fossil record of infections” and this ordering of spacers is important for the protection of the organism in which the CRISPR array is found. The implication contained within this hypothesis is that the leading spacer corresponds to the most recent attack, while downstream spacers match more ancient threats (11). In their analysis of the mechanism and physiological significance of type II-A polarised integration, McGinn and Marraffini showed that mutations in the LAS would lead to ectopic spacer integration. In this induced process, a sequence in the middle of an array was recognised as the anchor and consequently, the spacer was inserted on the downstream rather than on the leading side. Additionally, they found, under the same phage attack, a corresponding spacer located on the leading side (mimicking polarised spacer integration) conferred a fitness advantage to the host, compared to when the spacer was in the middle (mimicking ectopic spacer integration). In particular, they noticed that an ectopic integrated spacer can become the first spacer by deleting all segments between the mutated LAS and the new anchor (11). Spacer deletion and recombination between arrays have also been reported in other CRISPR loci (19). Additionally, Kupczok et al. (20) found, in an analysis of two γ -proteobacteria (type I) and two Streptococcus species (type II) that the evolution of spacer arrays is shaped more by deletion and acquisition than by recombination. This work would suggest that the ordering of spacers in CRISPR arrays might be quite dynamic and therefore potentially structured by natural selection and not just acquisition order.

**Figure 1.**
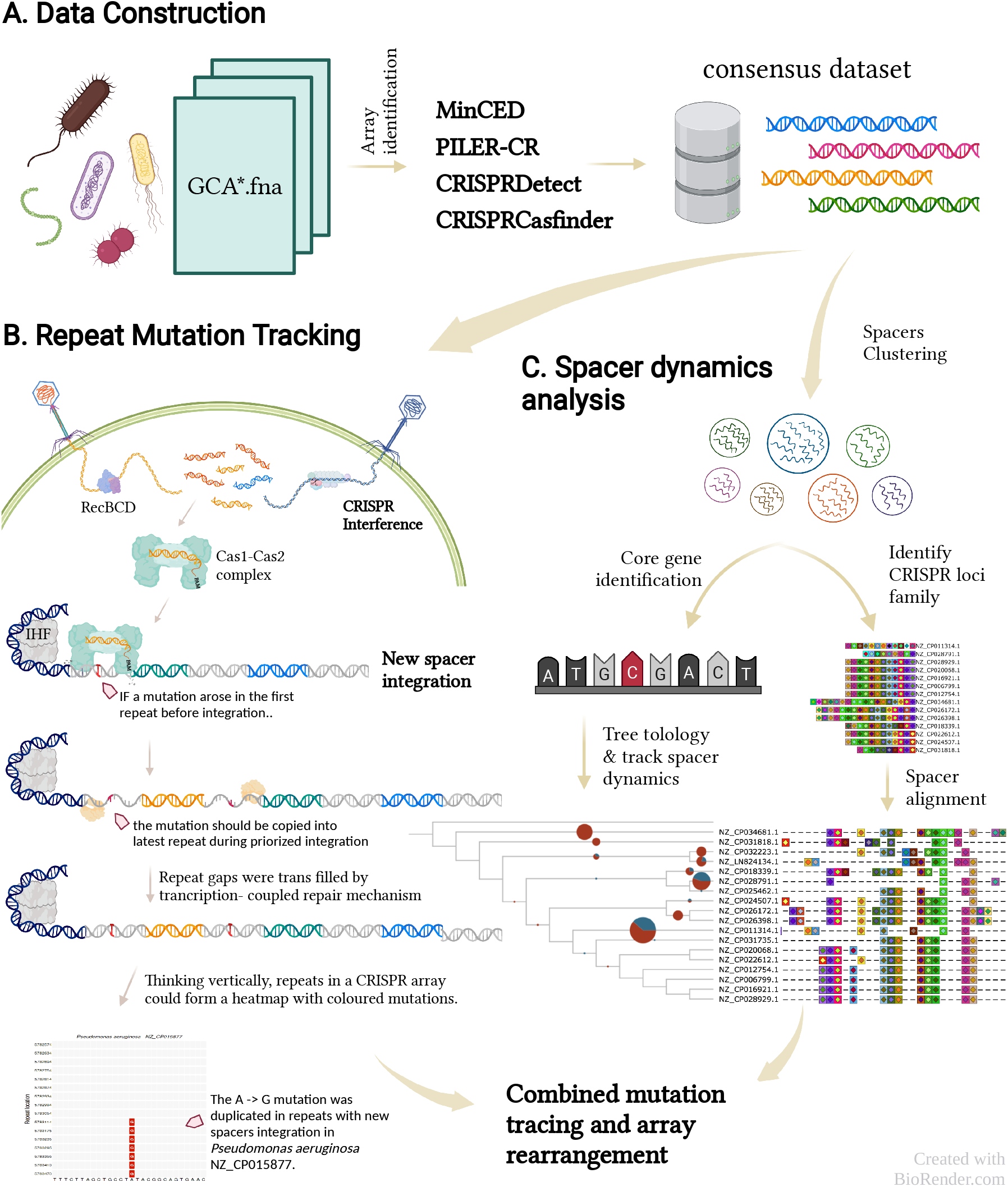
Workflow. After complete nucleotide genomes were downloaded from NCBI (A), CRISPR arrays were identified by four published programs and *cas* operon subtypes were classified by CRISPRCasTyper. An array that is identified by more than two programs is organised into the consensus dataset to carry out the following research: repeat mutation tracking (B) and spacer dynamics analysis (C). An array is comprised of direct repeat and spacer, and the evolutionary process could be visualised in these two approaches, respectively.

In this paper, we hypothesise that in dynamic and complex environments, natural selection would favour the active reordering of spacers in CRISPR loci by recombination and deletion, thereby potentially providing a boost in overall immunity against phage. In this scenario, spacer order might reflect fluctuating pressures from a dynamic external environment, rather than the chronological timeline of invasion. In addition, recombination between loci (20) and HGT across species (21, 22) can both increase spacer diversity and expand immunity range. In this study, we wished to investigate the frequency and extent of dynamic repeat re-ordering in CRISPR loci and how it competes with, and potentially overcomes the expected chronological ordering in CRISPR loci.

We compiled a dataset of 12,461 complete Archaeal and Bacterial genomes and identified their CRISPR-Cas loci. We used two different approaches to investigate spacer evolution. First, we traced spacer insertion and deletion in one CRISPR locus by tracking mutational patterns in adjacent duplicated repeats. This approach identifies changes within a single locus. We then focussed on the spacers themselves by combining phylogenetic trees with manually aligned CRISPR loci clusters to identify spacer gains, losses, and duplications across a diversity of clearly related CRISPR loci. These analyses show that CRISPR loci deviate substantially from the chronological-order-of-infection model and instead they manifest diverse histories, replete with dynamic patterns of insertion, deletion, recombination and ‘flip flopping’.

## Method and Materials

### Dataset Construction

A total of 277 Archaeal genomes and 12,184 Bacterial genomes were retrieved from the National Center for Biotechnology Information (NCBI) RefSeq database (23) in January 2019. Species information and download paths are available from https://github.com/JMcInerneyLab/CRISPRsharing/tree/master/dataset.

To compile a dataset of CRISPR arrays, we used a combination of MinCED (24), PILER-CR (25), CRISPRDetect (version 2.4) (26) and CRISPRCasfinder (27). The sizes of repeats are normally 23 to 47 bp (1) but extra-large repeats of 50 bp also have been identified (26). Repeats are nearly identical within CRISPR loci. Therefore, to find all possible CRISPR arrays, the lengths of repeats were set to between 23 to 55 bp during the execution of all programs. Descriptions and other parameters of these programs are listed in Supplementary S1. Considering the different programs return different results for CRISPR array identification, all arrays that were identified by at least three of the four programs were collated into a “consensus dataset” (Figure 1A). CRISPR-Cas subtypes were identified by CRISPRCasTyper (version 1.6.1) (28).

### Repeat Patterns

In order to track spacer evolution, we constructed alignments of the repeats that are found between every pair of adjacent spacers (Figure 1B). We represent the repeats in the alignments in the order in which they are observed in the array. In other words, the first repeat in an array is placed as the top sequence in the alignment and the last repeat is the bottom sequence. For the purposes of ordering the repeats from 5’ to 3’, we noted orientation as predicted by CRISPRDirection, from the CRISPRCasFinder and CRISPRDetect software suites. As outlined in Figure 2, any nucleotide found to be identical to the array’s consensus sequence is simply coloured grey. If a mutation has arisen, we have coloured the nucleotide according to the colours in the figure legend. In this way, new mutations can be easily observed. By representing the arrays in this way, we characterised the patterns of mutation into five groups.

**Figure 2.**
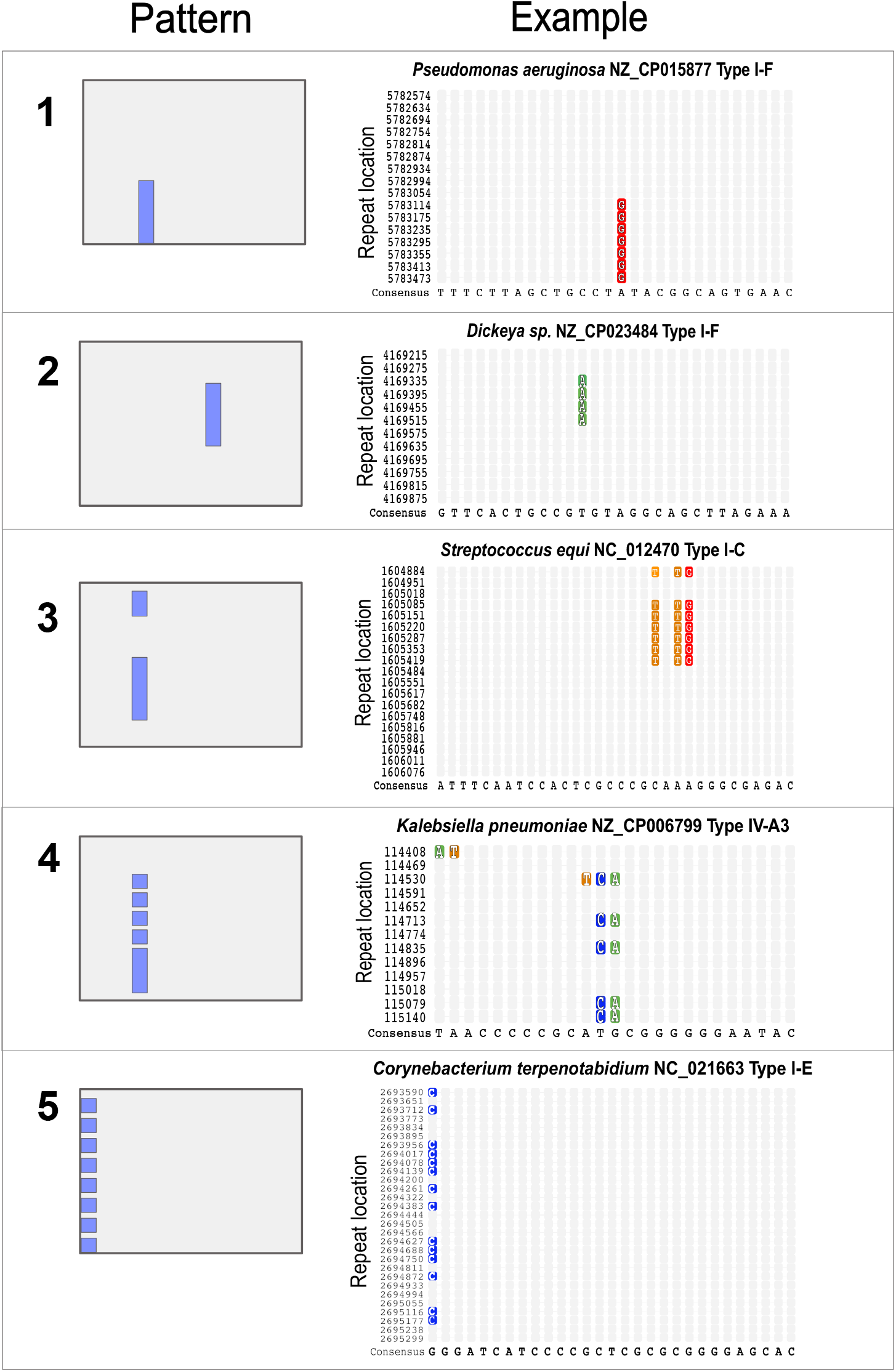
The different categories of repeat mutation patterns. On the left-hand side are schematic representations of the five different types of spacer patterns that we observed. On the right-hand side are specific examples of each pattern. The majority-rule consensus nucleotide is in grey, and the minor variant is outlined in colour. Cytosine is in blue, Thymine is in orange, Adenine is in green, and Guanine is in red. The spacers are ordered in the same order as in their respective arrays, with the first row of the alignment corresponding to the first spacer (at the 5’-end) in the array and the last row corresponding to the last spacer (at the 3’- end).

During dataset assembly, to minimise any potential influence of sequencing error, only CRISPR loci that had at least three repeats that contained a position that departs from the consensus were collected. In total, 1095 CRISPR loci were identified that have trackable mutated repeats and 1095 repeat maps were drawn using the ComplexHeatmap package (29) (data available at https://github.com/JMcInerneyLab/CRISPRsharing/tree/crispr/Repeats). In those cases where locus extension is sequential, unreversed and non-recombinant, we expect to observe a mutation pattern where a mutation arises in a repeat within the array, and persists through at least two, though possibly more, rounds of repeat duplication, right up to the beginning of the array. However, in our dataset, we observed a more diverse and complex picture, and the patterns we observed are summarised in Figure 2. Pattern 1 describes the aforementioned expected situation where a mutation arises in the repeat and this mutation is subsequently conserved through at least two duplications of the repeat. Pattern 2 involves a mutation arising and subsequently persisting through at least two duplication events but then there is ostensibly a reversal back to the original nucleotide character state. Pattern 3 depicts a more complex ‘flip flop’ pattern. A possible explanation of such a pattern might be that a mutation arose, a reversal of this mutation occurred, and then the mutation arose yet again. However, other explanations such as recombination could also result in such a pattern. Pattern 4 is when the ‘flip flop’ pattern is seen several times. Finally, we recognise Pattern 5, for mutations that only occur on the edge of the array.

### Spacer Patterns

In addition to the analysis of repeat mutations, we further investigated spacer evolution by tracing gains and losses throughout the evolutionary history of each CRISPR locus, using a combination of phylogenetic analysis and ancestral locus reconstructions.

Firstly, to group similar CRISPR arrays in the consensus dataset, we used the global alignment approach implemented in UCLUST (version 2.1), taking into consideration of the short size of spacers (mostly 27 to 45 bp) and repeats (mostly 23 to 47 bp). This identified arrays with sets of spacers and repeats with high similarity to one another. The sequence identity cut-off was set as 90% for spacer comparisons and 92% for repeat comparisons. To cluster CRISPR loci into ‘families’, we used the Markov Cluster (MCL) algorithm with an inflation value of 2.0, which generates clusters based on stochastic flows in graphs (30). Only pairs with more than two similar spacers were included in the analysis. Additionally, we only selected array families that contained at least 10 members.

To visualise each group of clustered CRISPR arrays, we used CRISPRStudio (version 1.0) and the spacer mismatch threshold was set as 4 (31). Output from the various CRISPR identification programs was reformatted using bespoke scripts in order to use as input to CRISPRStudio (Figure 1C). The scripts to perform these conversions were written in Python (version 3.5.3). In order to reconstruct the evolution of individual CRISPR loci, spacer arrays were manually aligned in order to track changes (data available at https://github.com/JMcInerneyLab/CRISPRsharing/tree/crispr/CRISPR_family).

To understand spacer dynamics in an array family, we constructed a “core gene” phylogenetic tree, *i*.*e*. using those genes found in every member of the group. In our analysis, the genomes in each array group always belonged to the same genus, therefore the outgroup for each phylogenetic tree was selected from a different genus, but the same taxonomic family. All genomes were annotated, using Prokka (version 1.13.7) (32), and these annotated genomes were used as input files for Roary (version 3.13) (33). Core genes were then aligned using MAFFT (version 7.453) (34) and a maximum likelihood gene tree was inferred using IQ-TREE (version 1.6.1) (35) using the GTR+I+G model. Confidence in phylogenetic hypotheses was evaluated using bootstrap resampling (1,000 replicates).

To analyse the gain and loss of spacers within each gene family, it is necessary to reconstruct the ancestral character states at each internal node in a phylogenetic tree (36, 37). We used the program Tree Analysis Using New Technology (TNT) (version 1.5) (38) in conjunction with spacer presence-absence (0/1) matrices and the core gene phylogenetic trees. The dynamics of spacer gains and losses were plotted onto the corresponding tree nodes and presented using the iTOL online tool (39).

## Results

Four programs (MinCED, PILER-CR, CRISPRDetect, CRISPRCasFinder) were used to identify CRISPR-Cas systems present in 277 archaeal and 12,184 bacterial complete nucleotide genomes from the NCBI RefSeq database. The results obtained from all four programs indicate a similar proportion of putative CRISPR arrays across all Archaea. However, the four programs produced quite different results when used to analyse bacterial genomes (Supplementary Table ST1). PILER-CR and CRISPRCasFinder identified more CRISPR loci in Bacteria compared with CRISPRDetect and MinCED. To conservatively compare and analyse CRISPR loci across species, a consensus dataset was constructed that only contained CRISPR loci that were identified by at least 3 programs. As shown in Supplementary Table ST2, 889 CRISPR loci were identified from 229 archaeal strains (82.7%) and 10,703 CRISPR loci were identified from 4,947 bacterial strains (40.6%) in the consensus dataset. The most common spacer length was 32 bp, followed by 36 bp, whereas the most common length of a direct repeat was 29 bp, followed by 36 bp (Supplementary Figure S1).

According to the output of CRISPRCasTyper, across the 11,592 putative CRISPR arrays, 6,230 arrays (54.7%) have recognisable *cas* genes in the adjacent neighbourhood (<= 10,000 bp window), 4,369 arrays (37.9%) have distant *cas* genes, while 966 arrays (8.3%) were found without adjacent *cas* genes (these are known as Orphan CRISPR arrays (40)). In addition, when analysing the arrays with adjacent *cas* operons, 2,529 CRISPR arrays were located downstream of *cas* genes, 2,139 upstream, and interestingly, 1,534 arrays were found to be situated between protein-coding *cas* genes and 19 arrays have overlapped with *cas* genes.

### Direct Repeat Tracking

We first depicted the evolution of spacers by tracking mutations in their proximal repeats. As noted previously, when a spacer adapts into a host genome, both the innate and primed procedures trigger the duplication of the first direct repeat (41). If a mutation occurs before duplication, this mutation is reproduced in the following repeats unless a reverse mutation occurs. From all putative CRISPR arrays in the consensus dataset, our filters returned 1,095 arrays that have trackable mutations in their repeat regions, and we plotted these mutations as a form of heatmap image. Although these trackable patterns account for only 9.5% of the whole consensus data, we expect that similar processes are taking place across the entire dataset, though they remain unobservable because no trackable mutations arose. All repeat mutation patterns are interactively available at https://shiny.its.manchester.ac.uk/mqbpryo2/RepeatM.

Pattern 1 (Figure 2) conforms to the current understanding of the spacer adaptation process. This pattern indicates that a mutation arose once as the CRISPR array was growing and this mutation was subsequently retained throughout further duplications. This pattern is observed in 495 (40.8%) CRISPR loci, meaning that it is the most common among all the patterns (Figure 3). Interestingly, however, we also observed many instances of patterns 2 (219 loci, 18.1%), 3 (248 loci, 20.4%), and 4 (165 loci, 13.6%), and together these amount to almost 30% more than the number of observations of pattern 1. Patterns 2 and 3 can arise either because mutations take place that subsequently reverses, or alternatively, they can arise because of insertions of spacers into the middle of the array. Pattern 4 is observed less often and this pattern probably results from mutational flip-flopping in the direct repeats or, alternatively, recombination within CRISPR arrays. The presence of non-canonical patterns 2, 3 and 4 demonstrate either frequent superimposed substitutions take place in a small number of sites, while most other sites remain mutation-free, or that recombination and middle spacer integration take place frequently, in addition to polarised insertion and deletion.

**Figure 3.**
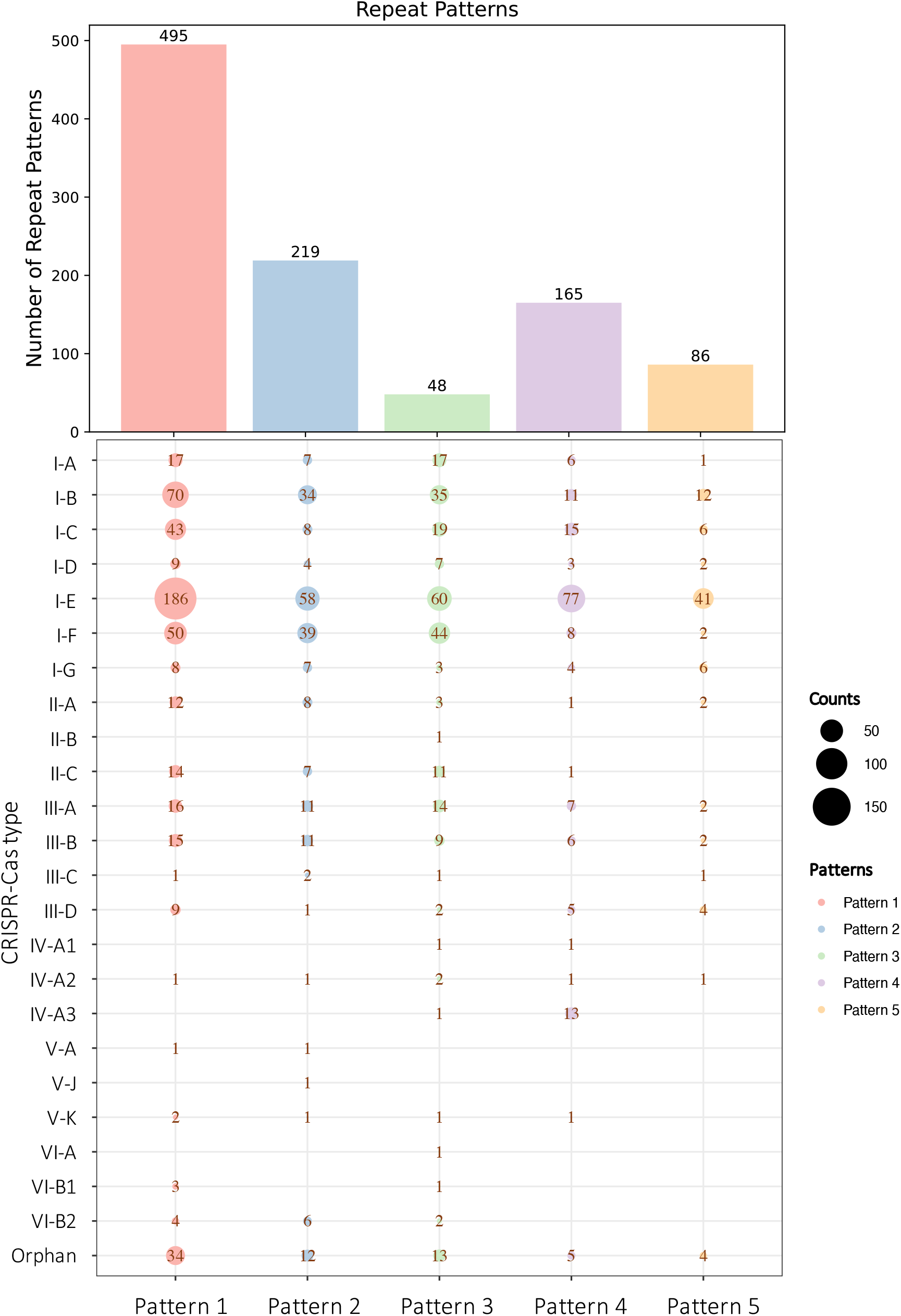
Numbers of observations of the patterns are outlined in Figure 2.

Finally, Pattern 5, which is present in the fewest loci (86 loci, 7.1%), could arise by either of two mechanisms. One, previously reported in the subtype I-E system (42), suggests that PAM can donate its last nucleotide during spacer integration. To investigate further, we identified the subtypes of all 1095 samples (Figure 3) based on results from CRISPRCasTyper *cas* clusters. We found that type I-E was present in 47.8% of all edge samples. The remaining samples which show the edge pattern may indicate the role that PAM played in this subtype. We note that we cannot rule out incorrect breakpoint identification between spacers and repeats. During identification, the first (or last) nucleotide of a spacer could be inadvertently included in the repeat sequence, a situation that reflects the difficulty with CRISPR locus delineation. Putatively homologous spacers in each array family are coloured by the same colour, and all loci were manually aligned (Figures 5, 6, and 7).

### Spacer Dynamics Analysis

Repeat-mutation trace analysis can only reflect dynamics within one array and the number of arrays that have recognisable mutations is limited. Consequently, we used a second approach to understanding array evolution. We identified “partially homologous CRISPR arrays” based on the similarity of their spacer content, and then analysed spacer evolutionary dynamics by reference to the phylogenetic tree uniting the genomes in which the partially homologous arrays were found. In our dataset, a total of 87 array families were found to contain 10 or more partially homologous array members (Figure 4). The program CRISPRStudio was used to visualise the dynamics of spacer changes between closely related arrays across genomes.

**Figure 4.**
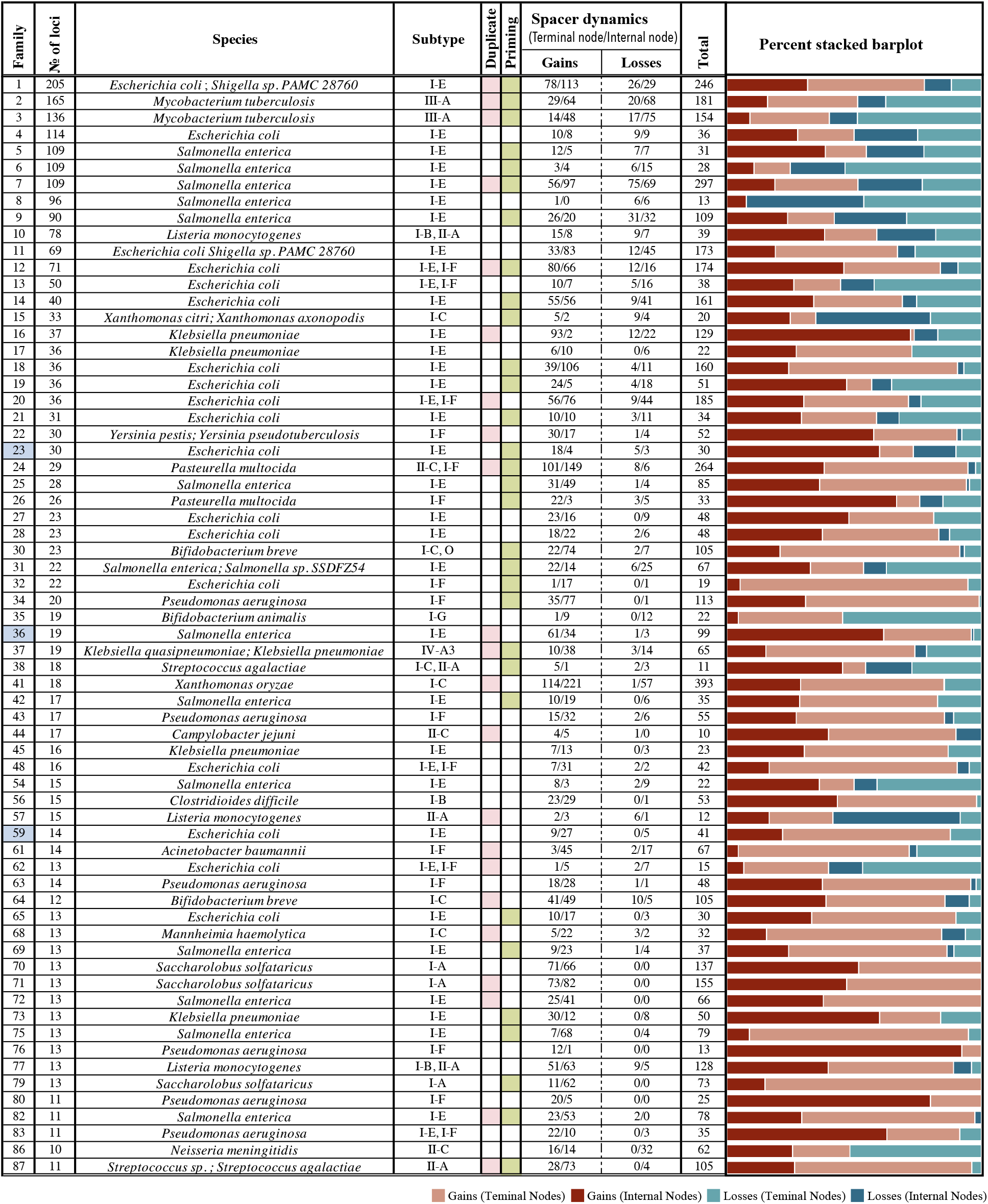
Spacer dynamics in 66 CRISPR array families. Details of spacer gain and loss events on tree nodes were summarised in the table and shown in percent stacked charts. 21 CRISPR array families that have less than 10 spacer activities (Supplementary Figure S2) were omitted in the table. Priming and duplication events were coloured by pink and green, respectively.

**Figure 5:**
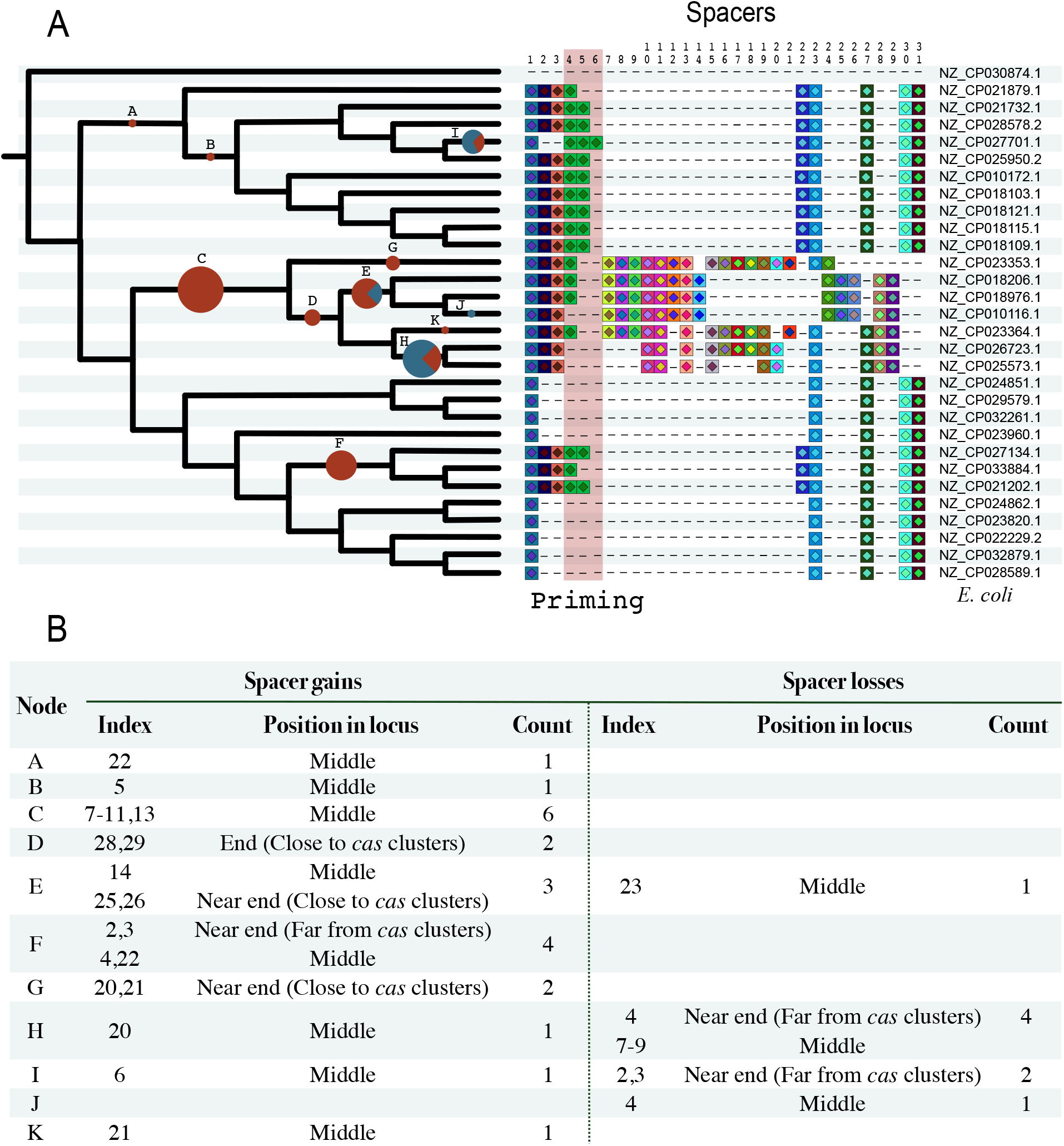
Aligned CRISPR arrays with the associated phylogeny for Family 23. (A) CRISPR array Family 23 contains 29 strains of *Escherichia coli* with *Raoultella sp. X13* used as an outgroup. The phylogenetic tree was inferred from core genes extracted from the respective genomes. The pie charts on the internal branches reflect the number of events hypothesised to have taken place on the branch, with the red segment indicating the proportion that are gains of spacers, while the blue segments reflect the proportion of events that were spacer losses. Spacer gain and loss details are organised in the below table (B). Spacer that locates within 4 repeats near terminal is regarded as near the end in this project (same as Figure 6).

**Figure 6.**
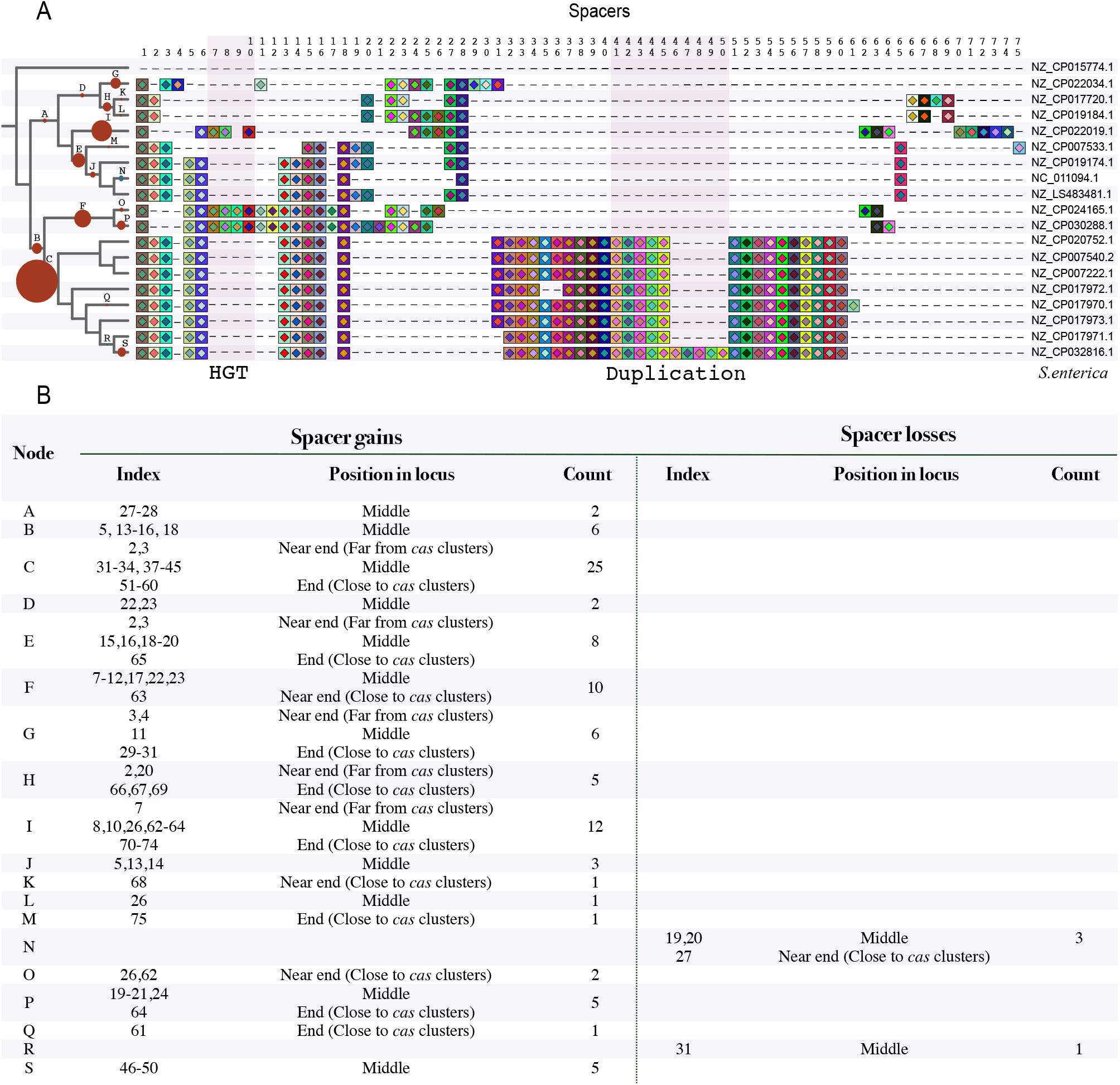
Aligned CRISPR arrays with the associated phylogeny for Family 36. (A) The phylogeny was constructed from core genes common to all strains. Gains and losses of spacers in strains and nodes are represented by the pie charts. Red represents gain and blue represents loss. The size of a pie chart is related to the total number of changes. Spacer gain and loss details are organised in the below table (B).

**Figure 7.**
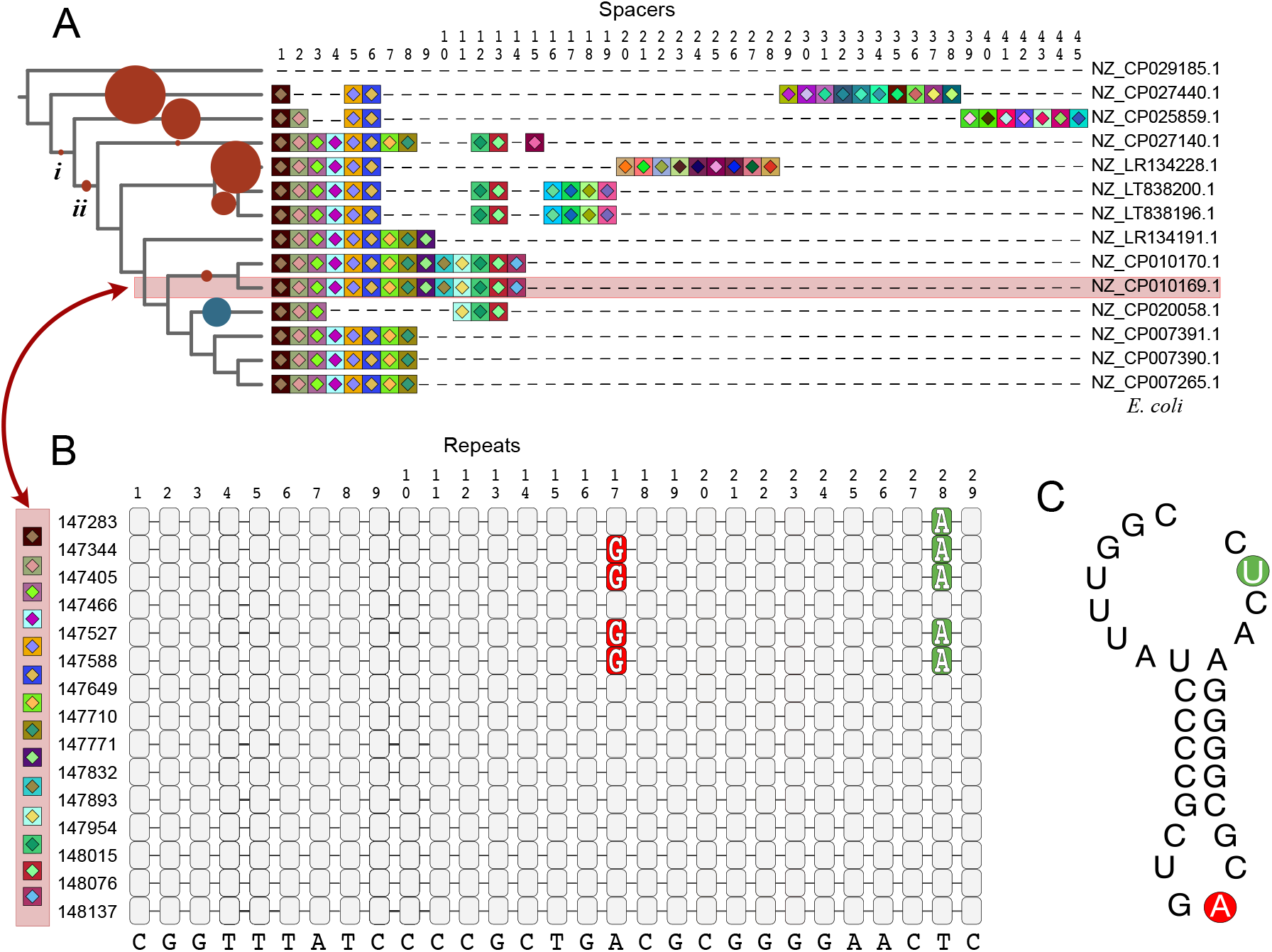
Arrangement of the Spacers and Repeats for CRISPR array Family 59 (a) with recognisable mutation patterns (b-e). (A): Phylogenetic tree of the backbone genes of the isolates, with aligned CRISPR arrays positioned at the tips of the tree. (B): The arrangement of the spacers with the minority mutations highlighted. (C): A generic RNA folding diagram of the repeats, with the mutations highlighted.

In our hands, almost all array families could be well aligned, except for two families that have extra-long arrays and aligning these long arrays exceeded our manual processing ability. Therefore, in any given family, spacers shared between loci were typically in the same order, though with interesting exceptions. Since we ignored all spacer order information while clustering, extensive recombination within CRISPR arrays would have made the process of alignment impossible. Thus, the fact that we could align CRISPR spacers in 85 families suggested that the rate of recombination within a locus during spacer evolution is rare. One thing that needs to be noted, however, is that our choice of cut-offs when using the UCLUST algorithm has undoubtedly influenced the arrays that we have compared. We are comparing arrays with extensive overlap and, therefore, conservation. More stringent spacer mismatch settings can often link arrays together with few shared spacers. A denser sampling of pangenomes has the potential to change our perception of how often recombination occurs. Using a 90% spacer similarity cut-off, we find that recombination is relatively rare, but this picture could change when tolerating more changes.

Here, we present two examples (Figures 5 and 6) from the 87 array families that met our criteria. We select these two examples because the array lengths are appropriate to show in one figure and because they exemplify specific evolutionary processes, such as priming and duplication. Spacers in the manually aligned CRISPR loci families were marked with integers.

CRISPR array Family 23 (Figure 5) contains 29 strains of *Escherichia coli* with *Raoultella sp. X13 was* used as an outgroup. Gains and losses of spacers occurred independently in different lineages multiple times, though, in this dataset, it appears that most changes are relatively recent, especially gain events (Figure 4). Evidence for the gain of new spacers supports the possibility that there are other methods of integration apart from polarised integration. For example, spacers 14, 25 and 26 in strains NZ_CP018206.1 and NZ_CP018976.1, NZ_CP010116.1 are in the middle of their respective arrays, but their restricted phylogenetic distribution suggests that they were inserted on a later branch than spacers 28 and 29 on the leading side. Curiously, spacer 24 is found in these three genomes and also in a more distantly related genome. While the insertion of 14, 25 and 26 at the stem-lineage subtending NZ_CP018206.1 and NZ_CP018976.1, NZ_CP010116.1 is the most parsimonious explanation for the observed pattern, it does not rule out more complex, unobserved events of gain and loss. Also, a series of potential priming events (spacers 4-6 in the red square of Figure 5) can be tracked through CRISPR alignment. Evidence of priming was detected in 33 out of 87 samples, a situation that has recently been confirmed as being relatively common (Figure 4) (15).

Another example of a complex pattern of spacer gains and losses is found in CRISPR array Family 36 (Figure 6) which includes 18 strains of *Salmonella enterica*, with *Lelliottia amnigena* being used as an outgroup. Spacers 4 and 11 in NZ_CP022034.1 were integrated recently but located downstream of spacers 22 and 23 that were integrated earlier in the history of the genomes, providing additional evidence for non-chronological spacer integration. Spacers 7, 8, and 10 are shared between two distant genomes NZ_CP022019.1 and NZ_CP024165.1 (or NZ_CP030288.1). It is unlikely that two perfectly identical sections of an array have arisen independently one after another. What is more likely is that a single three-spacer segment was constructed in one genome and subsequently transferred to another. Moreover, as shown at the bottom of Figure 6, a five-spacer gain pattern (spacers 46-50) detected in NZ_CP032816.1 indicates an independent array duplication. Duplications were detected in 23 out of 87 samples (Figure 4) (20). Overall, different evolutionary processes including polarised integration (including priming), ectopic integration, duplication, loss, and HGT are reported, which have cooperatively led to an intricate spacer history.

### Combined mutation tracing and array rearrangement

Among our 87 examples, only one family – CRISPR array Family 59 (Figure 7) - has the two properties of having alignable arrays, and having recognisable mutations in direct repeats and these repeats are adjacent to spacer evolution events. The repeats are 29 bp in length and two positions have undergone mutations. Figure 7C depicts the most stable RNA secondary structure as inferred by RNAfold (43) and the two sites with mutations are outlined. These two positions are both on loops of the RNA structure and do not bond with one another.

If we follow the pattern of spacer acquisition, the most parsimonious scenario is that spacer 1 was acquired first, then spacer 2 – their distribution across the tree and their position in the arrays would both point to the same inference. The distribution of spacers 5 and 6 also implies that they are quite ancient and their origin tracks to an internal branch (*i*). At the base of this clade, position 17 in the repeat array was probably an A nucleotide and position 28 was also an A nucleotide (though we obviously cannot rule out other alternatives), although position 28 was coloured as a mutation in Figure 7B. This is because we used the conservative repeat as a module rather than the first repeat. At position 17, the first mutation, an A-to-G, was introduced when this first repeat was duplicated (Figure 7B, S3).

The fourth repeat (position 147466 in the genome, Figure 7B), which is located between spacers 3 and 4, has the same sequence as the repeats 7 (147649) to 15 (148137). This presents a puzzle for which the answer is not clear. Either two mutations appeared simultaneously in repeat 4 and were immediately back-mutated when a new spacer was acquired, or this repeat was actually constructed later in the evolution of this clade of organisms, but recombination has moved it to position 4 in our dataset. Several scenarios could fit this pattern, but the scenario that the repeats and spacers are in chronological order, is not very parsimonious, requiring multiple simultaneous mutations in the same positions.

Together, these results indicate that CRISPR array evolution is not simply inserting in a chronological way and deleting under selective pressure, but a much more complex process along with unanswered mechanisms.

## Discussion

As with many datasets, we cannot rule out the possibility that the subset of data in our analyses is in some way biased. We applied conservative filtration steps to obtain alignable arrays and identifiable mutations in repeats. The most likely bias we have introduced is one that will underestimate the true amount of change that occurs within arrays.

The patterns that we outline and provide examples for in Figure 2, particularly patterns 2, 3 and 4 could quite conceivably arise by inter- and intra-genome recombination. A section of a CRISPR array, with near-identical repeat regions, could quite easily become integrated into an array in a different genome, either by transduction, plasmid-mediated transfer, or following the death of a cell and release of the DNA into the environment, transformation. If we combine the information from the repeat analysis and the spacer analysis, this seems to be a common scenario. We can clearly see in Figures 5 and 6 that there are transfers of portions of CRISPR arrays between genomes. Therefore, the relative contributions of within-array and between-array recombination it is still an open question. However, if the patterns we observe in this relatively small sample are indeed an indicator of the situation globally, then CRISPR array recombination takes place on a massive scale worldwide every day.

Although this study does not directly address the issue of the balance between mutation, drift, and selection, this is a key element to the puzzle. There is evidence that a positive fitness effect is associated with having a spacer near the beginning of a CRISPR array (44). However, it must be said that our observations of spacer shuffling are relatively rare when we consider the rigorous spacer order in loci across 85 CRISPR families. Therefore, we need to consider the likely causes. First, direct repeats are substrates for the recombination mechanisms of any cell, so the recombinations we observe could be a side-effect of having such a system in the cell. This is the side-effect hypothesis. The alternative hypothesis is that these recombination events are observed because of natural selection favouring recombinations that bring the “most important” spacers to the top of the array. However, aligned groups of array families suggest the rigorous order of CRISPR loci was rarely affected by recombination within a locus. Nevertheless, recombination can also occur between loci on one host genome or across different organisms. During recombination, when a new spacer is acquired from other loci, the original spacer with repeat will be deleted (20, 45). Another possible explanation is that a spacer is introduced into the array and it is immediately recombined so that any subsequent events make it look like there is no recombination on the spacer. In addition, duplication of spacers, as a result of recombination (19, 20, 46), was identified in 26.4% of array families. By contrast, the study from Kupczok et al. (20) suggested that CRISPR evolution was mainly shaped by acquisition and pervasive deletion instead of recombination.

The selection model for prokaryotic pangenomes (47–49) suggests that selection would strongly affect the order of spacers in these arrays – favouring certain genome variants that have spacers near the front of the array that are in immediate need, or that are in need of the most often. The rapid change that we see happening in CRISPR arrays reflects the ever-changing nature of natural ecosystems.

We propose that activities such as recombination or novel integration mechanisms can both cause this pattern. It has been reported that mutation in conserved LAS can also result in middle spacer acquisition in type II-A CRISPR-Cas systems, which is termed “ectopic spacer integration”. Strains that lack LAS, or have a mutant-LAS can potentially use another sequence within a CRISPR locus as an anchor during integration to guarantee precise positioning (11). Although through ectopic spacer integration, mutations in repeats should still be preserved, the patterns we observe open the possibility of other undiscovered mechanisms by which spacers might integrate.

Although spacer turnover is rapid in CRISPR loci, a special pattern of spacer retention was observed. Spacers located on the edge of an array were highly conserved across different array families, like an anchor. A similar model was identified in another study, that called this phenomenon “trailer end clonality” (50, 51). However, the study from McGinn and Marraffini (11) showed that the immunity conferred by a spacer in a CRISPR locus is related to their position in that locus. Newly integrated spacers are the most important for the cell when it is attacked. Also, spacers located at the very beginning have been shown to produce more transcribed crRNA (46, 52–54) and lead to more robust immunity. For example, crRNA from Position 1 is produced at a twofold higher level than from Position 5 (11). Overall, the location of spacers is very important to a CRISPR locus. The fitness effects of having spacers in different positions on a locus in a complex, changing ecosystem, remain to be studied.

