## Supplemental Files for "High Frequency of Dynamic Rearrangements In Crispr loci"

### Supplementary Files

**S1. Brief Introduction of CRISPR identify programs and** **the execute Setups**

There are four programs (MinCED, PILER-CR, CRISPRDetect and CRISPRCasFinder) were used to detect CRISPR-Cas systems from species in our dataset. I will briefly describe them here, as well as the ways in which they were used in order to identify CRISPR arrays.

***MinCED***

MinCED (version 0.2.1) is short for Mining CRISPRs in Environmental Datasets, a Java-based program that derives from program CRT (21). It is designed for detecting CRISPR repeats and spacers from complete genomes or environmental databases.

***PILER-CR***

PILER-CR (version 1.06) is a C++ based program that can be used to identify CRISPR arrays rapidly and accurately (22). On top of detecting CRISPR arrays, PILER-CR can also cluster similar direct repeats into groups with the help of software MUSCLE (46), in which the minimum identity is set as 75%. The cutoff for repeat length was set to the same values as was used in MinCED.

***CRISPRDetect***

CRISPRDetect (version 2.4) (23) is a part of the CRISPRSuite, a CRISPR detection, viewing, analysis, and comparison package. It supports both web and command platforms, and input can be fasta, gff or gbk files. CRISPRDetect uses the CRISPRDirection algorithm (also in the package CRISPRSuite) to identify CRISPR array direction (47). In particular, CRISPRDetect has a scoring system that evaluates arrays based on nine known biological properties such as repeat length, similarity to referenced sequences, *cas* genes and so forth. The detailed calculation can be found in Table S1. The developers have suggested that an array with a score above 4.0 can be graded as good. However, CRISPRDetect can only predict *cas* genes from gbk formatted input rather than the fasta format that was used in this study. Therefore, 3.0 was set as a conservative quality score cutoff during the execution of the program. In contrast to other programs, the threshold of spacer lengths is not fixed during execution, cutoffs change with repeat length of the array.

***CRISPRCasFinder***

CRISPRCasFinder (version 4.2.17), an update of the web program CRISPRFinder, is one of the newest programs designed to predict CRISPR arrays (24). It is similar to CRISPRDetect, CRISPRCasFinder seeks to verify the direction of the CRISPR array (using CRISPRDirection). CRISPRCasFinder also has a scoring system that ranks four evidence levels based on the number and conservative level of repeats and spacers. Detailed level rules are listed in Table S2. Any potential CRISPR array that satisfies the two criteria that the evidence level is higher than level 2 and also it has a known direct repeat sequence, is retained. In particular, CRISPRCasFinder can call the dependency program Macromolecular System Finder (MacSyFinder) during execution to predict Cas proteins. Then the identified cluster Cas proteins are used to classify system subtypes (48).

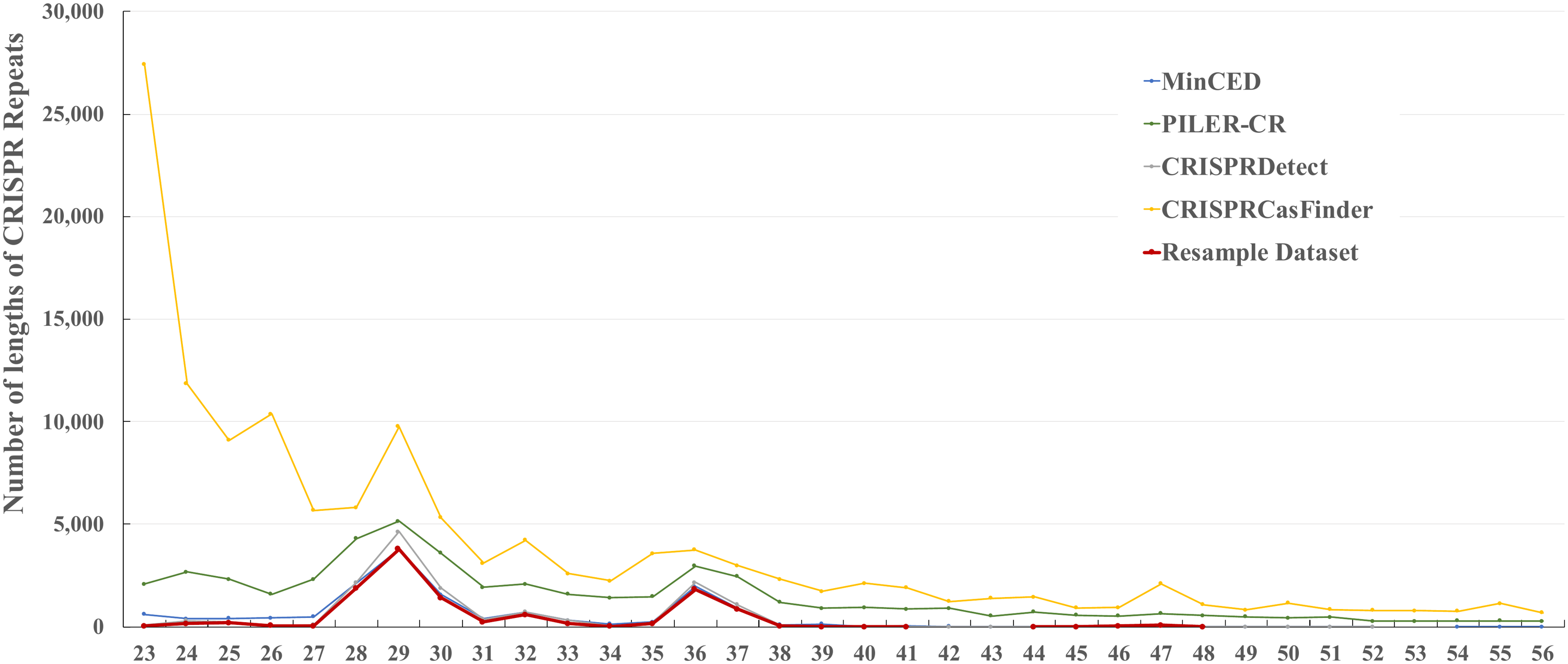

(a)

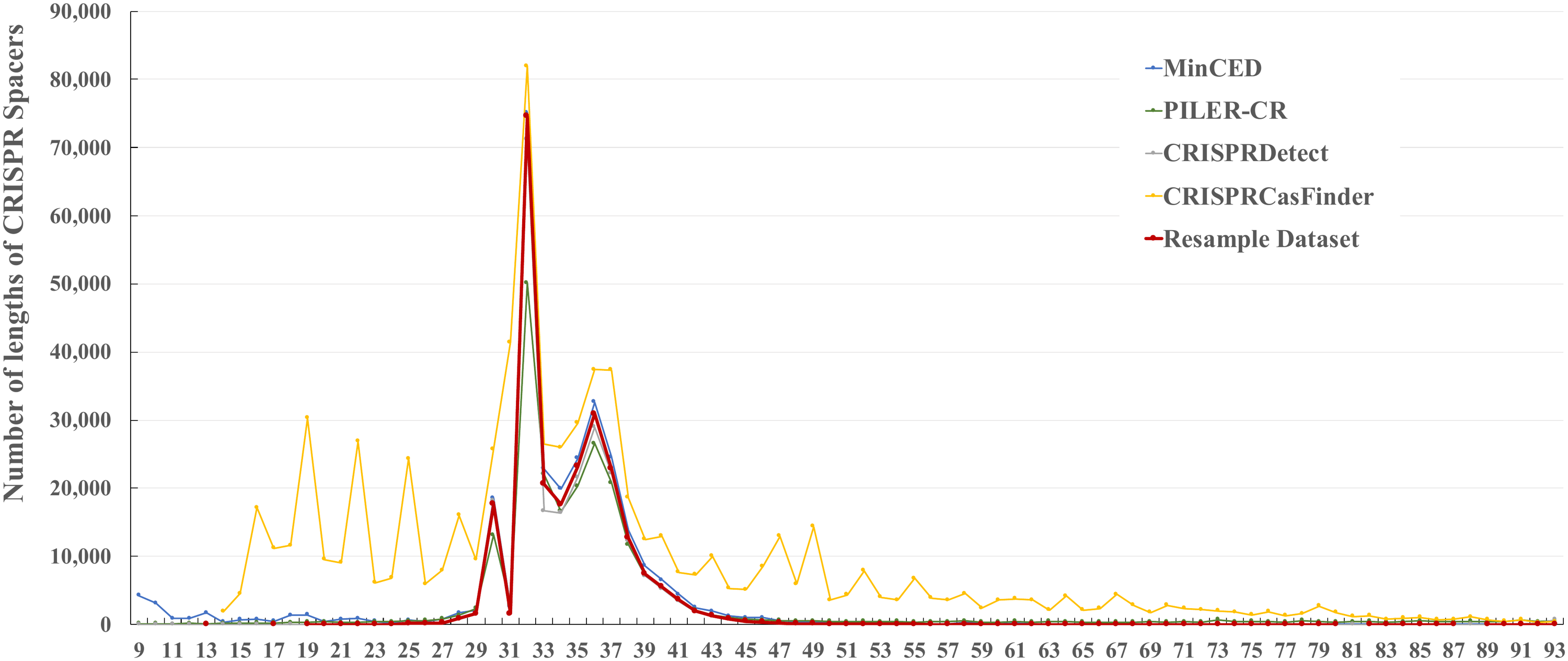

(b)

**Figure S1. Numbers of CRISPR repeats (a) and spacers (b) that were identified by program MinCED, PILER-CR, CRISPRDetect, CRISPRCasFinder and found in the consensus dataset.**

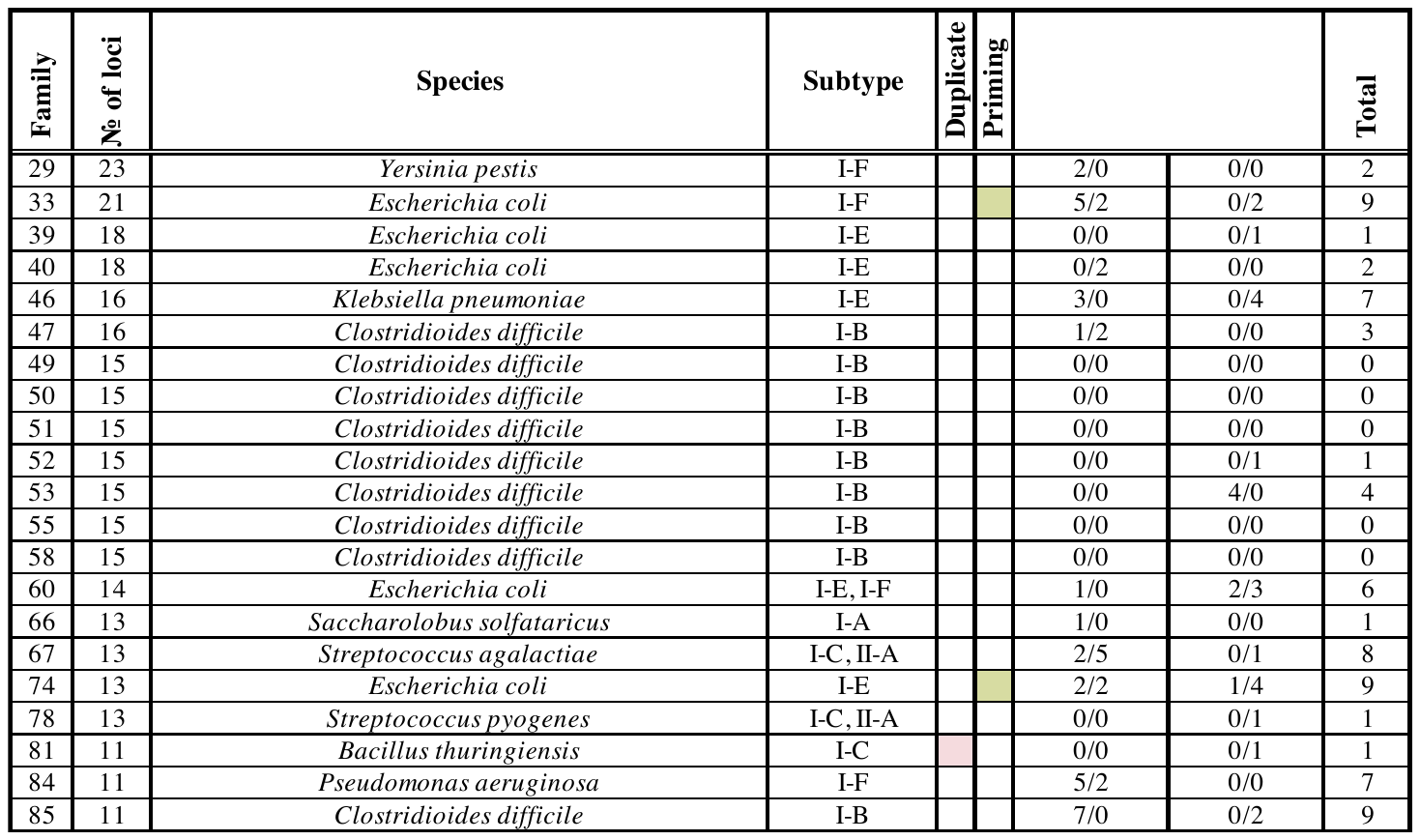

**Figure S2. Spacer dynamics in 21 CRISPR array families** have less than 10 spacer activities. Priming and duplication events were colour by pink and green, respectively.

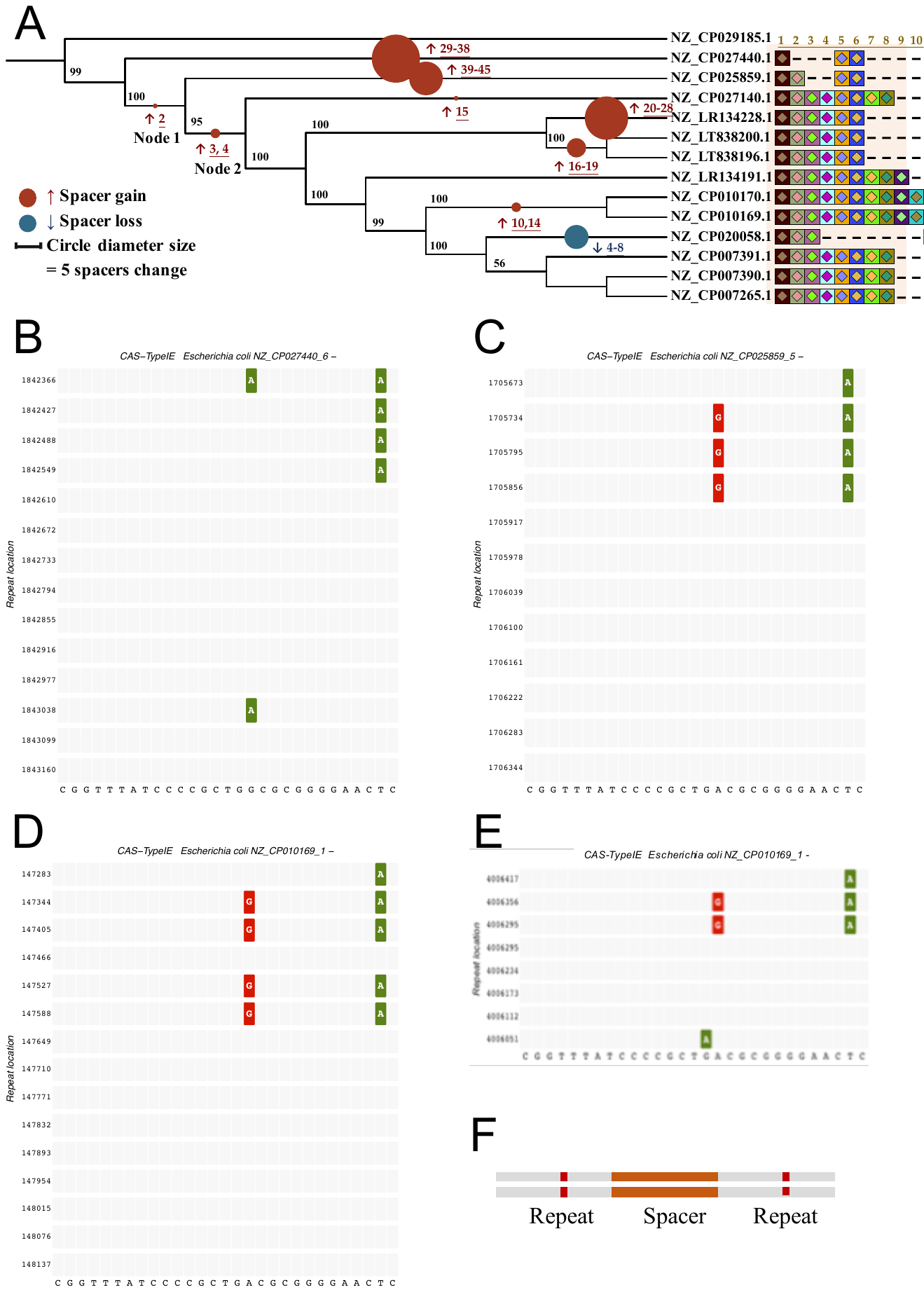

**Figure S3. CRISPR array alignment of Family 59 (A) with recognisable mutation patterns (B-E).** All loci in this family are in the reverse orientation, which means spacer 1 which is located at the 5’end is the oldest spacer. Also, since CRISPR arrays follow a “repeat-spacer-repeat” pattern (Figure S3F), a spacer is located between two repeats. According to the canonical model, a repeat, as depicted in our alignment, could have served as the template for the repeat below it. This might or might not be the case, since insertions, deletions and rearrangements within the array might have taken place in the interim period. In Family 59, the deepest diverging genome that contains this CRISPR family is NZ_CP027440.1 (Figure S3B). In this CRISPR array we see the ‘expected’ pattern of mutation – a T to C mutation arose in the second-last nucleotide of the spacer and this mutation was copied three times, as evidenced by its presence in four flanking repeats. As we move along the phylogenetic tree, we see the insertion of spacer 2 on the branch subtending Node 1, the node is ancestral to all genomes on the tree except for NZ_CP027440.1 and the outgroup. As shown Figure S3C, in NZ_CP025859.1, site A mutated to G and conserved to the following integration of spacer 5. However, abnormal repeat duplications (Pattern 3) were identified in Node 2 while spacers 3 and 4 integrated (as shown in NZ_CP010169.1, Figure S3D). The repeat that is next to spacer 4 still followed previous mutations while spacer 3, located upstream next to the locus, conserved the repeat. Considering that the evolutionary history of spacer 5 was missed, we proposed three possible conjectures that may explain this pattern. First, spacer 3 initially integrated downstream of spacer 5 but was shuffled to the middle through recombination. Second, a replacing recombination may happen between loci who share the same repeat; this could result in deletion of the repeat with mutation and insertion of the original conserved repeat. The third is probably due to ectopic spacer integration with an unknown repeat duplication mechanism. In the most recent strain NZ_CP020058.1, there were 5 spacer losses (spacers 4 to 8), which only left repeats with two spacers in Figure S3E. Intriguingly, we found spacers 1, 5, 6 are shared across all loci in Family 59 in the same order but integrated with different repeats, which may result from HGT or indicate a robust fitness benefit of these three spacers

**Supplementary Table ST1. Numbers of species with identified CRISPR arrays in Archaea and Bacteria using four programs.**

| MinCED | 239 (86.3%) | 6,079 (49.9%) |
| --- | --- | --- |
| PILER-CR | 259 (93.5%) | 9,804 (80.5%) |
| CRISPRDetect | 237 (85.6%) | 5,399 (44.3%) |
| CRISPRCasFinder | 252 (91.0%) | 9,718 (79.8%) |

**Supplementary Table ST3. CRISPR statistics for the consensus dataset.**

|  | Archaea | Bacteria |
| --- | --- | --- |
| Genomes that have CRISPR loci | 229 (**82.7%)** | 4,947 (**40.6%)** |
| Genomes that have identified *cas* clusters | 212 (76.5%) | 4,430 (36.4%) |
| CRISPR loci in plasmids | 11 (4.0%) | 187 (1.5%) |
| *cas* clusters in plasmids | 8 (2.9%) | 151 (1.2%) |
| Total number of CRISPR loci | 889 | 10,703 |
| Range of spacer number in a locus | 2 to 245 | 2 to 587 |
